# Spatial transcriptomics defines the mechanisms of hiPSC-derived stem cell-mediated repair in human articular cartilage

**DOI:** 10.64898/2026.07.22.740000

**Authors:** Yolande F.M. Ramos, S. Sana Sayedipour, Georgina Shaw, Margo Tuerlings, Timo Schomann, Helena E.D. Suchiman, Davy Cats, Frank Barry, Rachid Mahdad, Hailiang Mei, Luis J. Cruz, Mary Murphy, Ingrid Meulenbelt

## Abstract

We here determined therapeutic efficacy and mode-of-action of human induced pluripotent-derived therapeutic stem cells (hiMSCs) across *in vivo* mouse and *ex vivo* human osteoarthritis models. hiMSC treatment in DMM-mice significantly reduced OARSI damage scores, which was affirmed by a decrease in the catabolic marker Mmp13 and an increase in the anabolic marker Col2. These treatment effects appeared, irrespective of modifying factors such as xeno-free media or thermosensitive hydrogel carrier. Subsequently treatment of hiMSC+gel in human osteoarthritic cartilage explants showed a transcriptome-wide significant activation of the cholesterol and sterol synthesis pathways marked by genes such as *MVD, DHCR7, MSMO1, FABP3*. Additionally, we showed that these changes alleviated OA-associated imbalances of the cellular Zinc-ion homeostasis pathways, represented by genes such as *MT1F, MT1G, MT1H and SLC30A1*. Spatial transcriptomics then sensitively captured that hiMSC+gel treatment evoked, specifically at the superficial cartilage layer, a consistent upregulation of healthy chondrocyte markers such as *CHAD, ACAN, FRZB*, and *SOX9*, alongside a suppression of catabolic and inflammatory mediators such as *SERPINE1, SPP1, MMP13, ADAMTS5*. Our findings link therapeutic outcomes of hiMSC treatment to precise spatially resolved molecular changes in human tissue, that would otherwise be obscured by heterogeneous cell populations. Collectively our study highlighted that hiPSC-derived stem cell therapy (hiMSCs) could provide a scalable ‘off-the-shelf’ solution to treat osteoarthritis, with strong prospects for clinical applications in the near future.

## INTRODUCTION

Osteoarthritis (OA) is a painful and debilitating progressive joint disease characterized by cartilage degradation, formation of osteophytes and subchondral sclerosis leading to limitations of joint function. The impact on society and health care is high, with around 595 million people affected worldwide and a projected increase between 50% and 95% by 2050 dependent on joint type.(*1*) Despite the burden on society and health care systems disease-modifying therapy is still limited.(*2*) In recent years, however, human autologous mesenchymal stem cells (hMSCs) derived from different sources such as bone marrow and adipose tissue were tested as advanced therapy medicinal product (ATMP) for OA. Given the positive effects on pain, function and symptoms in hMSC-treated patients, stem cell therapy was proposed as a potential regenerative treatment in OA.(*3*) Despite these promising results, wide clinical application of autologous hMSCs is still limited due to the invasive collection procedures and restricted possibilities for *ex vivo* expansion.(*4-6*) Specifically, expansion of hMSCs is known to evoke changes in therapeutic potency due to phenomena such as replicative senescence and associated metabolic changes.(*7, 8*) Moreover, there is intrinsic heterogeneity in the potency of collected autologous hMSCs that is difficult to assess prior to the clinical application.(*9*) Together these issues have hampered clinical application of hMSCs as potential novel disease modifying treatment of OA. To overcome these limitations, there has been an increasing focus on the generating therapeutic stem cells derived from human induced pluripotent stem cells (hiPSCs).(*10*) For that matter, we have generated and characterized mesenchymal stem cells derived from hiPSCs (hiMSCs). We showed that hiMSCs closely resemble autologous hMSCs in terms of morphology, immunophenotype, and trilineage differentiation potential.(*11*) Moreover, hiMSCs allowed for prolonged expansion in the absence of senescence-related impairments.(*11-13*) Even more, it allows for selection of the most potent therapeutic cells prior to cryopreservation and cell banking at large scale.

To determine therapeutic efficacy and mode-of-action of hiMSCs, we here applied a multi-tiered experimental approach. First, we studied chondroprotective effects of hiMSCs *in vivo* in a destabilization of the medial meniscus (DMM) mouse model of OA. This model reliably induces cartilage degradation, synovial inflammation, and subchondral bone changes comparable to those observed in OA patients. Herein, we directly compared the efficacy of hiMSCs with conventional bone marrow derived hMSCs (hBMSCs) while utilizing xeno-free medium and a recently developed thermosensitive hydrogel for intra-articular (i.a.) injection(*14*) as potentially modifying factors. To confirm translational relevance of chondroprotective features observed, we subsequently used hydrogel delivered hiMSCs to study therapeutic effect in *ex vivo* human OA cartilage explant model, and procured in-depth insight into the direct therapeutic mode-of-action on articular cartilage by applying spatial transcriptomics.

## MATERIAL AND METHODS

### Ethics approval

Ethical approval for the generation of hiPSCs from skin fibroblasts of healthy donors was obtained by the Medical Ethical Committee of the LUMC and is available under number P13.080. The control hiPSC line used in the current study was generated by the LUMC iPSC core facility from male skin fibroblasts (LUMC0004iCTRL10 (004) registered at the Human pluripotent stem cell registry. Cells were characterized according to pluripotent potential and spontaneous differentiation capacity by the iPSC core facility.(*15*) hBMSCs were derived from bone marrow aspirates of a healthy donor with approval from the NUI Galway Research Ethical and Galway University Hospitals Clinical Research Ethics Committees. Collection of cartilage explants from OA patients undergoing total joint replacement surgery is approved by the Medical Ethical Committee of the LUMC within the ongoing RAAK study(*16*) and available under numbers P08.239 and P19.013.

### Patient and Public Involvement statement

Patients or members of the public were not involved in the design, conduct, reporting, or dissemination plans of this research. However, the OA-research group at the LUMC involves OA patients in the research through regular meetings. To this end, the ‘Patient Participatie Artrose – Leiden’(PPA-Leiden) was founded in 2017. They meet regularly and their involvement entails, among others, proof-reading of patient information forms (e.g. the RAAK study).

### Cell culture

The hiMSCs applied in this study were generated previously from LUMC0004iCTRL10 cells using Stem Cell Technologies Mesenchymal Progenitor Kit following the manufacturers’ instructions with small modifications as described.(*17*) The hiMSCs were extensively characterized,(*11*) cryopreserved until use, and applied at passage 8. Likewise, applied bone marrow derived hBMSCs were previously collected, characterized, and cryopreserved until use at passage 4.(*11*) Cells were cultured either in serum-containing medium (DMEM-GlutaMAX medium (Gibco) supplemented with 10% fetal calf serum (Sigma-Aldrich), 100U/mL penicillin-100mg/mL streptomycin mixture (GiBCo), and 5ng/mL basic FGF (Peprotech)) or in PS medium(*18*)). The cells were maintained at 37°C with 20% O2 and 5% CO2, and passaged upon reaching around 80% confluence.

### Thermosensitive hydrogel

Synthesis and characterization of the hydrogel have been described in detail recently.(*14*) In short, a thermosensitive hydrogel delivery system based on poloxamer 407 (P407) with a self-assembling peptide (Palmitoyl-WKGNNQQNYQQ) were maintained at 4°C until use to avoid solidification. Hydrogel alone or mixed with cells was applied *in vitro* by pipetting or *in vivo* using a 30G needle. A temperature shift to 37°C results in gelation of the hydrogel which ensures its stabilization and localization over time for at least 60-72hr.(*19*)

### In vivo osteoarthritis model

Twelve-week old male C57BL/6J mice were purchased from Charles River Laboratories (Charles River, Chatillon-sur-Chalaronne, France). Animal procedures were all conducted at the Leiden University Medical Center and were approved by the Animal Welfare Committee (IvD) under number AVD1160020171405-PE.18.101.005. All mice were housed in groups in polypropylene cages on a 12hr light/dark cycle with unrestricted access to standard mouse food and water. Knee OA was induced in the mice by surgical destabilization of the medial meniscus (DMM) on the right knee joint (N=42) as described previously.(*20*) Further details of the *in vivo* experiment are described in the **Supplementary Materials and Methods** and experimental groups are shown in **Data file S1**.

### Human ex vivo osteoarthritis model

To determine the direct effect of stem cells on cartilage, in the absence of interference of any additional factors such as synovium, a previously developed *ex vivo* explant model with knee joints included in the Research in Articular Osteoarthritis Cartilage (RAAK) study(*16*) was applied. In total, nine donors were included (characteristics shown in **Data file S2**). Following generation of explants from the lesioned regions of the donor material (8 mm diameter) they were allowed to equilibrate in xeno-free medium for 72hr before the explants underwent a four-day treatment with 5×10^3^ hiMSCs cells with hydrogel (hiMSC+gel). An advantage of hydrogel usage is the ease to apply the treatment locally with cells remaining better in place while use of lesioned cartilage in the *ex vivo* model more authentically reflects aged OA cartilage as it occurs in the population.

### Correlation gene expression and Mankin score

Spearman correlations were calculated between Mankin scores(*21*) and the expression levels of all genes expressed in articular cartilage (n=20,048 genes and n=34 paired preserved and lesioned cartilage samples) from the RAAK study as determined before with bulk RNA-sequencing (RNA-seq).(*22*) Correlation coefficients were calculated in R statistical language using the Hmisc package and *P*-values were adjusted for multiple testing using the Benjamini-Hochberg false discovery rate (FDR) correction while considering FDR<0.05 to be significant.

### Histology and immunohistochemistry

Histological and histochemical analyses of joint tissues was performed according to our established protocols and have been described previously.(*19, 23*) In short, samples were fixed in 4% paraformaldehyde for 24hr, embedded in paraffin, and sectioned at 5 μm. Mouse joints including bone tissues were decalcified prior to embedding by applying a 5-day incubation step in Mol-Decalcifier (Milestone) at 37°C. Paraffin blocks (both mouse joints and human explants) were sectioned and stained with Hematoxylin & Eosin (H&E), with Safranin O/Fast green, or with toluidine blue (TB; Sigma-Aldrich) and mounted with Pertex (Sigma-Aldrich) as described before, and imaged with Zeiss Axioscan Z1 slide scanner. Both, antibodies for detection of COL2A1 (ab34712, Abcam, Cambridge, MA) and for MMP13 (sc-515284, Santa Cruz Biotechnology) were diluted 1:200 in PBS-5% BSA with an incubation overnight at 4°C. TNFRSF11B/OPG was detected with antibody EPR3592 (Abcam/Epitomics, Cambridge, MA), diluted 1:100 in PBS-5% BSA and also incubated overnight at 4°C. More details can be found in the **Supplementary Materials and Methods**.

### Assessment of histology and immunohistochemistry

For histological assessment of of the cartilage damage of the femur and tibia in knee joint, 20x magnifications were used for evaluation according to OARSI guidelines.(*20*) Scoring was performed by three independent researchers blinded to the conditions and to the scores of the other investigators. Results were averaged and used as representative score of the knee joint as described elsewhere.(*24, 25*) Quantification of IHC staining intensity for Col2 and Mmp13 were done using ImageJ-based image analysis. Microscopy images were processed by splitting the color channels, applying a rolling ball algorithm to remove background noise and segment the stained tissue area as described previously.(*26*) Scores for TNFRSF11B/OPG and COL2 expression in explants were determined as done previously in our group,(*27*) and based on consensus of two experienced researchers blinded to the samples.

### RNA-sequencing

RNA was extracted from cartilage explants using the RNeasy Plus Kit (Qiagen) according to the manufacturer’s indications. Subsequently, RNA integrity values were determined with a 2100 Bioanalyzer (Agilent) which were at least 6. RNA sequencing (polyA enriched) was performed using the Illumina NOVAseq 6000 for control samples (n=7 preserved, n=12 lesioned OA cartilage samples from N=9 donors) and treated samples (n=17 lesioned OA cartilage samples from N=9 donors treated with hiMSCs in the thermosensitive hydrogel) according to the standard operating procedures based on the Illumina protocol for Paired-End Sequencing (Per sample ∼6Gb, 20 million Paired-End reads RNA-seq mapping). Quality control (QC) and downstream analyses of generated data were performed using the open-source BioWDL RNA-seq pipeline developed by the SASC team as described previously.(*22*) Further details are described in the **Supplementary Materials and Methods**. Enrichment for biological processes was determined with online available tool DAVID while including FDR significant genes.

### Spatial transcriptomics

Following RNA-seq readout of the hiMSC treatment of lesioned OA cartilage explants, an optimal, tailored, probe panel was designed with the available online Xenium Panel Design Tool (**Data file S3**) for genes to be detected at single cell level in their *in situ* geographic location (spatial transcriptomics). Spatial Transcriptomics was performed for two paired OA lesioned cartilage explants with or without hiMSC+hydrogel treatment. Target probes (N=295 genes) were selected with online available Xenium Panel Design Tool based on: genes that are FDR significantly differentially expressed upon hiMSC+hydrogel treatment as determined by RNA-seq; previously identified FDR significantly differentially expressed genes with OA pathophysiology and markers for cartilage damage correlating to Mankin scores.(*22*) Genes with extremely high expression levels in cartilage were excluded to avoid interference with detection of other (lower) expressed genes. With this we could compare changes upon treatment and also identify specific cell types or cell clusters with specific changes. Spatial Transcriptomic analysis on 5 µm sections of paraffin-embedded cartilage explants (n=2 paired OA lesioned cartilage explants from N=2 donors) was performed by the Leiden Genome Technology Center (LGTC) using the Xenium platform (v1; 10X-Genomics) according to the manufacturer’s protocol and initial data exploration was performed with online available Xenium Explorer 2.0.0 from 10X-Genomics. Data preprocessing and quality assessment were conducted using Squidpy.(*28*) Cells containing fewer than 10 detected transcripts were excluded, and genes expressed in fewer than five cells were removed from downstream analyses. Dimensionality reduction and clustering were performed using Uniform Manifold Approximation and Projection (UMAP) together with the Leiden algorithm.(*29*)

### Statistical analysis

OARSI scores were analyzed by applying linear generalized estimating equation (GEE(*30*)) using IBM SPSS statistics 25 to effectively adjust for dependencies among DMM experiments: OARSI score ∼ Hydrogel + cell source + (1|DMM+surgeon) and *P*-values were considered significant if they were smaller than 0.05. For RNA sequencing genes were considered differentially expressed (DEGs) upon treatment in comparison to controls if FDR was smaller than 0.05.

For cross-donor integration of spatial transcriptomic data, a fixed-effects meta-analysis was performed using METAL.(*31*) Donor-specific logFC and FDR values were used as input, and instead of simply averaging the input, METAL computes combined Z-scores with meta-FDR values to quantify the overall direction and significance of gene regulation across donors, with the Z-score indicating how many standard deviations a data point is from the mean.

## RESULTS

### Therapeutic potential of hiMSCs in comparison to hBMSCs in vivo using a DMM OA mouse model

**Figure 1A** shows a schematic overview of our experimental strategy to determine therapeutic efficacy of hiPSC-derived mesenchymal stromal cells (hiMSCs) in an *in vivo* OA mouse model, with destabilization of the medial meniscus (DMM) to induce OA related damage. Throughout, we investigated the therapeutic potential of hiMSCs in parallel with hBMSCs. Additionally, the effect of modifying factors such as our recently developed hydrogel delivery system(*14*) and type of medium (serum-containing or xeno-free PurStem (PS) medium(*18*)) were explored. To this end, three weeks after DMM surgery, mice received a single i.a. injection with different combinations of cells and media with or without hydrogel, randomly assigned to each mouse (**Data file S1**). The combinations hence experimental groups allowed us to study the independent chondroprotective effect of hiMSCs, hydrogel, and media parameters. Noteworthy is that hBMSCs were always applied together with the hydrogel (hBMSC+gel), therefore, we could not assess the effect of hBMSCs alone.

**Fig. 1.**
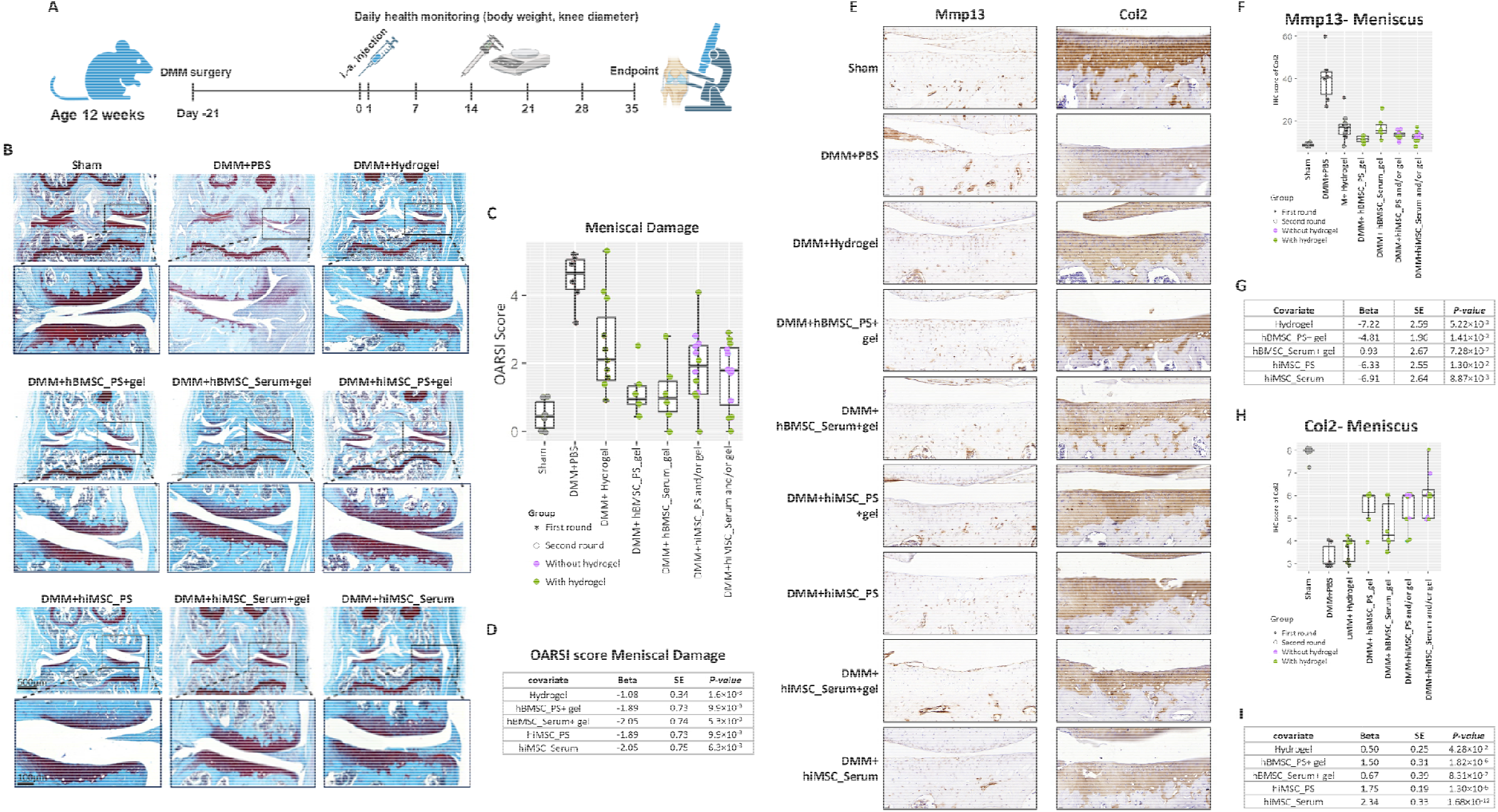
Therapeutic effect of hiMSC, hBMSC and hydrogel in DMM mice model. **(A)** Schematic outline of timelines in vivo experiment. Intra-articula (i.a.) injectior of experimental groups was performed 3 weeks after DMM surgery and samples were collected for analyses 5 weeks later. **(B)** Representative image of Safranin O/fast green staining for knee joint at 8 weeks post-surgery in the different experimental groups as indicated. **(C)** Boxplot showing the OARSI score of the meniscus lesions graded on a scale of 0 to 6 according to the OARSI scoring system and **(D)** GEE analyses of OARSI scoring indicating the independent therapeutic effect of hiMSC, hBMSC and hydrogel in the meniscus lesions. **(E)** Representative image of IHC for Mmp13 and Col2 for knee joint at 6 weeks post-surgery in the different experimental groups as indicated. Boxplots for quantitative analysis corresponding to Mmp13 **(F)** or Col2 (**H**) IHC (values determined by ImageJ software) and GEE analysis for Mmp13 **(G)** or Col2 (I) expression levels. All data are shown as means ± standard deviations (n=6; GEE: generalized estimating equations; First/Second time: first or second surgery round; IHC: Immunohistochemistry).

First, we compared cartilage integrity between the experimental groups, using the OARSI damage scoring system. As shown in **Figure 1B** negative control mice (Sham) had uniform Safranin O-staining and intact cartilage tissue that sharply contrasted with the unstained subchondral bone, reflecting minimal cartilage damage in the knee (average OARSI score: 0.3). On the other hand, the OA control mice (DMM+PBS) had markedly reduced Safranin-O staining of the cartilage with fibrillation and fissures to the bone reflecting severe cartilage loss and damage (average OARSI score: 4.5). Notably, the DMM mice treated with hiMSC and/or gel or hBMSC+gel exhibited intermediate Safranin-O staining and cartilage characteristics as compared to the control mice, with average reduction in OARSI damage score up to 1.2 (**Fig. 1C**). Moreover, cell expansion medium (serum-containing or PS) did not affect therapeutic efficacy of either hBMSCs or hiMSCs, reinforcing the translational suitability of clinically compliant conditions. To assess the independent effects on cartilage integrity of the treatment groups multivariate regression analyses was performed. As shown in **Figure 1D** OARSI damage scores of the hiMSC and hBMSC+gel groups were significantly reduced relative to untreated controls (DMM+PBS), with similar effect sizes (Beta ∼ -2). Additionally, we showed that OARSI damage score in the hydrogel group also showed a significant independent reduction, albeit with a more modest effect size (Beta=-1).

We next evaluated the effects of treatment on knee cartilage through immunohistochemistry (IHC) for matrix metalloprotease 13 (Mmp13), a catabolic marker for cartilage degradation, and collagen type 2 (Col2), an anabolic marker for cartilage integrity (**Fig. 1E-I**). Six weeks after i.a. injection, negative control mice (Sham+PBS), showed no Mmp13 expression and high Col2 expression levels (**Fig. 1E** and **H**). Vice versa, untreated DMM mice (DMM+PBS), showed high Mmp13 and low Col2 protein expression. The DMM mice treated with hiMSC and/or gel or hBMSC+gel exhibited a beneficial decrease in Mmp13 expression and an increase in Col2 expression relative to the control mice (DMM+PBS) although the magnitude of response varied between the experimental groups (**Fig. 1G** and **1I**).

Upon assessing the independent beneficial effect of the treatment parameters by multivariate regression analysis, we showed for Mmp13 that particularly the hydrogel and hiMSC mice groups had a strong and significant beneficial reduction (Beta<-6.3, *P*<1×10^-2^), which outperformed the hBMSC+gel group (Beta ∼ - 4.8, **Fig. 1F-G**). On the other hand, for Col2, we showed that particularly the stem cell treatment groups (hBMSC+gel and hiMSC) had a pronounced and consistent anabolic Col2 upregulation (Beta>1.5, *P*<1×10^-6^) that outperformed the independent effect of the hydrogel (Beta=0.5, *P*=4×10^-2^, **Fig. 1H-I**). Notably, the effect of hiMSC treatment groups was not affected by cell expansion medium (serum-containing or PS), albeit that particularly the hMBSC cultured in serum did not show any beneficial effects. Taken together, our results showed that hiMSCs and/or hydrogel showed a dual independent chondroprotective effect as evidenced by a reduction in the OARSI damage score, a reduction in the catabolic Mmp13 protein expression, and an increase in the anabolic Col2 expression.

### Introduction of the human ex vivo OA lesioned cartilage explant model

Based on the beneficial findings in mice, we next aimed to study the therapeutic effect of hydrogel-delivered hiMSCs in a human context, hereafter referred to as hiMSCs+gel treatment. In doing so, we used lesioned human *ex vivo* cartilage explants (macroscopically affected cartilage with average Mankin score of 6.8) obtained from patients (N=9, donors) that underwent joint replacement surgery due to knee OA (RAAK-study(*16*)). Donor characteristics are summarized in **Data file S2**). Human lesioned cartilage explant were subsequently treated with hydrogel-delivered hiMSCs for 5 days (**Fig. 2A**). This allowed us to study the treatment effect of hiMSCs+gel on cartilage integrity and chondrocyte phenotypes, without the interference of additional tissues such as the synovium. To determine therapeutic efficacy and mode-of-action of hiMSCs+gel, we compared histology, bulk RNA-seq and spatial transcriptomics between lesioned untreated and lesioned treated cartilage explants. Preserved (macroscopically normal cartilage explants, average Mankin-score of 4.9) were used as reference. Prior to establishing the treatment effect of hiMSC+gel across these output measures, we aimed to select a sensitive marker that could serve as a proxy for structural articular cartilage damage as assessed by Mankin score. To select such a marker, we analyzed human articular cartilage samples (N=34) for which both bulk RNA-seq and Mankin-scoring was previously obtained.(*22*) This showed a strong and consistent positive correlation between *TNFRSF11B*, encoding osteoprotegerin (OPG) protein expression and Mankin score (ρ=0.81, *P*=8.6×10^-9^, **Fig. S1, Data file S4**).

**Fig. 2.**
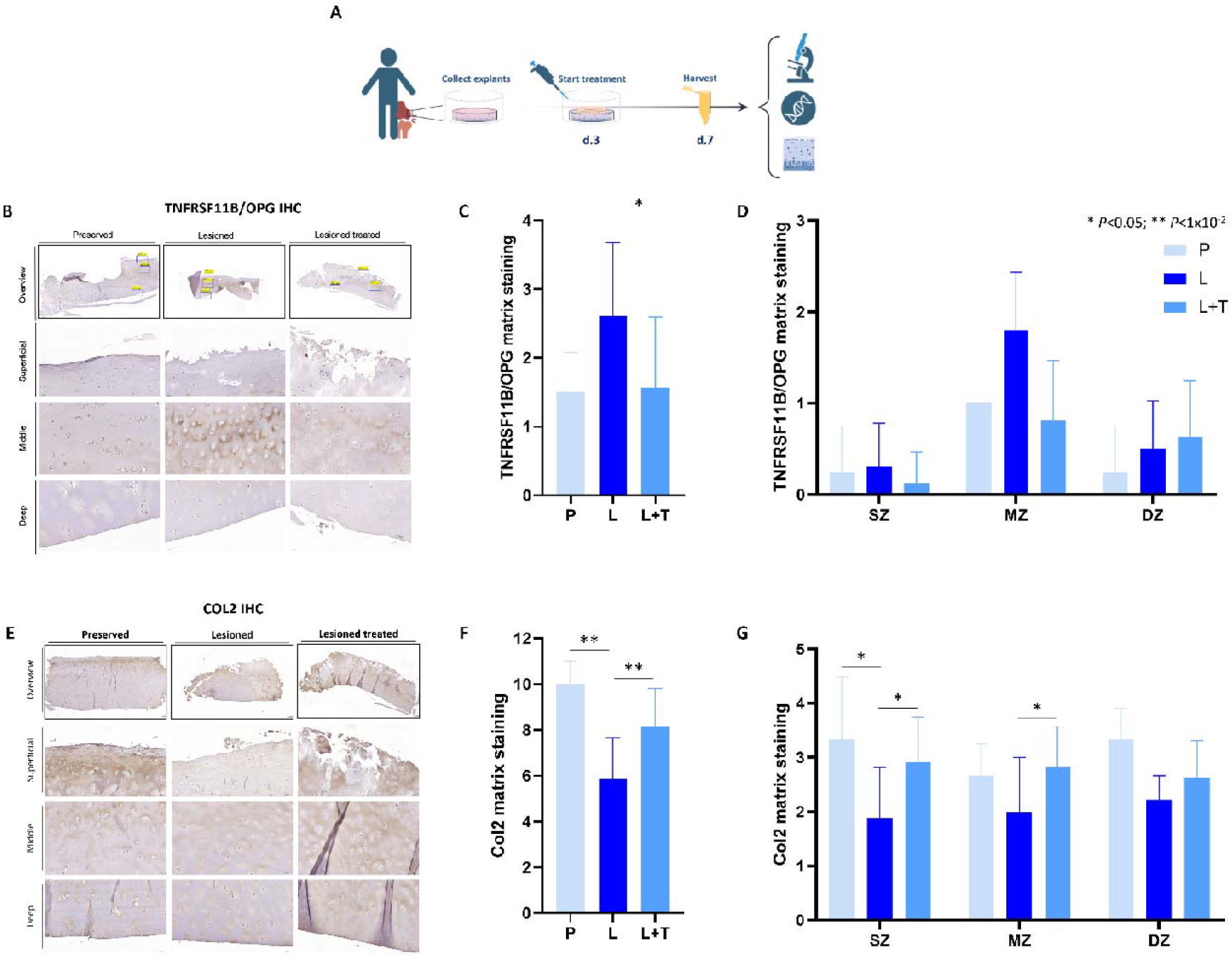
Treatment of lesioned explants with hiMSC+gel improves cartilage phenotype as assessed with immunohistochemistry. **(A)** Schematic outline of *in vitro* model. Human *ex vivo* cartilage explants are collected (ongoing RAAK study) and three days later treated with hiMSCs+gel for harvest and analyses at day 7. **(B)** Representative IHC images for expression of *TNFRSF11B* encoding osteoprotegerin (OPG) in preserved, lesioned, and lesioned treated human *ex vivo* cartilage explants across the superficial, middle, and deep zone of articular cartilage. **(C)** Quantified scoring for TNFRSF11B/OPG: total score and differentiated scores in superficial, middle and deep zone of cartilage. **(D)** Representative IHC images for expression of COL2 in preserved, lesioned, and lesioned treated human *ex vivo* cartilage explants across the superficial, middle, and deep zone of articular cartilage. **(E)** Quantified scoring for COL2: total score and differentiated scores in superficial, middle and deep zone of cartilage (\**P*≤0.05; IHC: Immunohistochemistry; P: preserved; L: lesioned; L+T: lesioned treated (hiMSC+gel); SZ: superficial Zone; MD: Middle Zone; DZ: Deep Zone).

### Effects of hiMSC treatment on cartilage integrity of lesioned human OA explants

To assess the effect of hiMSC+gel effect on the integrity of lesioned OA cartilage explants, we performed immunohistochemistry (IHC) for OPG encoded by *TNFRSF11B* and performed semi-quantitative blinded scoring that was depicted for the overall as well as for the zonal staining intensity. As shown in **Figure 2B**, hiMSC+gel treatment resulted in a significant lower overall expression of OPG protein (**Fig. 2C-D**). Moreover, zonal semi-quantitative scoring demonstrated that this effect was most pronounced in the middle zone of lesioned cartilage. Next, we analyzed the hiMSC+gel treatment effect on the extracellular matrix protein collagen type 2 (COL2) in a similar way. As shown in **Figure 2E** hiMSC+gel treatment affirmed an overall significant higher COL2 protein expression, particularly in the superficial and middle zone of lesioned cartilage as determined with semi-quantitative blinded scoring (**Fig. 2F-G**). Together, these data show that hiMSC+gel treatment of human lesioned cartilage explants showed a direct beneficial effect on the integrity of the extracellular matrix, with zonal differences.

### Transcriptome wide effects of hiMSC treatment of lesioned human OA explants

To gain further insight into the molecular processes underlying hiMSC+gel treatment and to characterize transcriptome-wide changes in the osteoarthritic chondrocyte phenotype, we next performed bulk RNA-seq of human preserved, lesioned (untreated) and lesioned hiMSC+gel treated cartilage explants. As shown in **Figure 3A-B** (**Data file S5**), hiMSC+gel treatment resulted in 61 FDR-significant differentially expressed genes with 48 upregulated and 13 downregulated genes. Notable highly significant upregulated genes included *MVD* (FC=2.7, FDR=8.50×10^-18^), *MSMO1 (*FC=2.8, FDR=9.6×10^-18^), *HMGCS1* (FC=3.1, FDR=1.8×10^-16^), *DHCR7* (FC=2.9, FDR=2.0×10^-11^), *ACAT2* (FC=2.7, FDR=3.0×10^-15^) and downregulated gene *ABCA1* (FC=-3.0, FDR=4.0×10^-10^). Subsequently, gene enrichment analysis for biological process highlighted that these genes are particularly linked to cholesterol biosynthetic processes (FDR=1.2×10^-33^; **Data file S6**). Additionally, we found that promotors of these genes were significantly enriched for transcription factor binding sites (TFBSs) recognized by Nuclear Factor Y (NF-Y), a highly conserved trimeric transcription factor complex that binds specifically to CCAAT-box sequences (**Data file S6**).

**Fig. 3.**
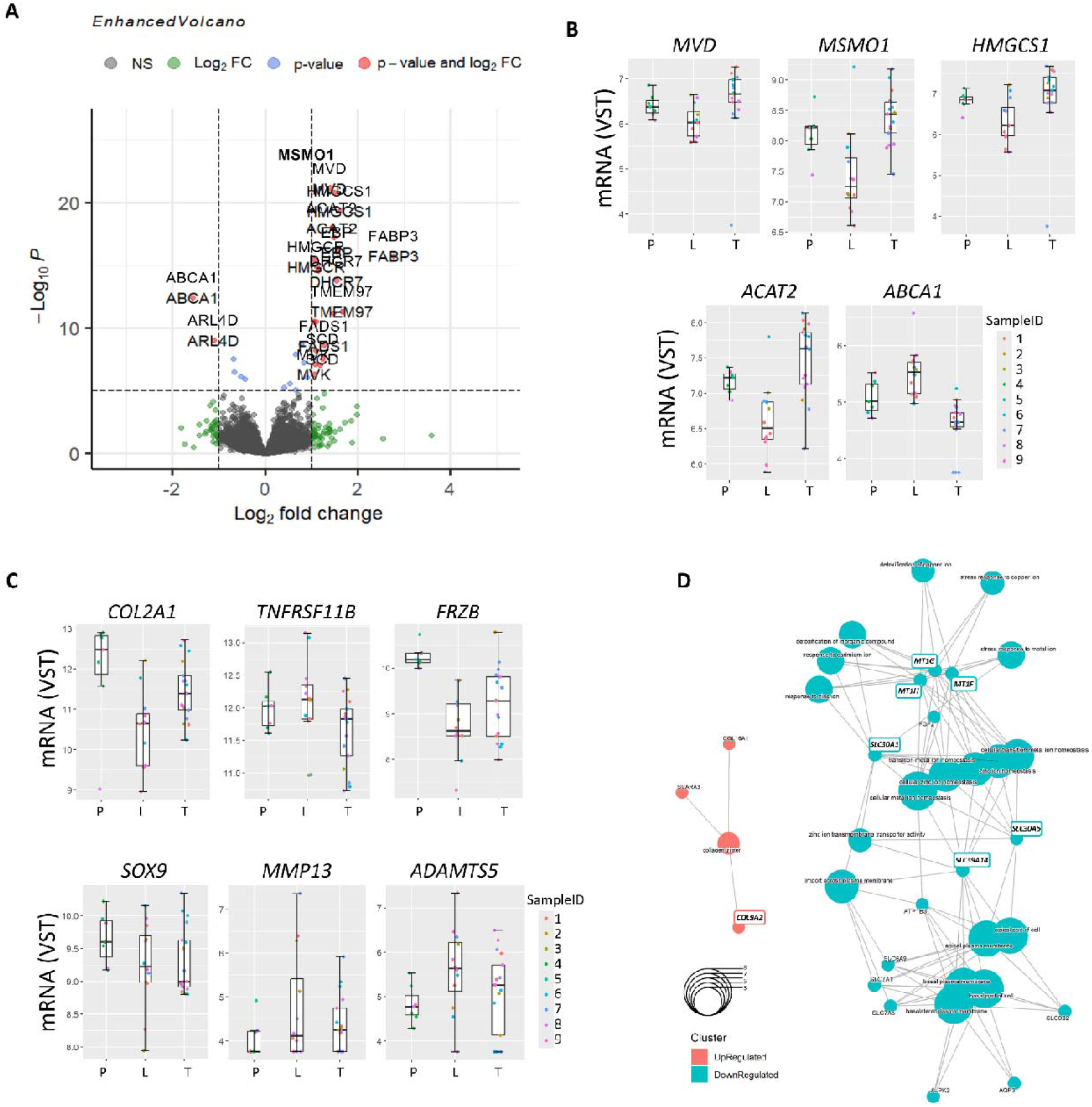
Gene expression profiles in preserved, lesioned, and hiMSC+gel–treated cartilage explants. **(A)** Volcano plot of differentially expressed genes from bulk RNA-seq comparing hiMSC+gel–treated (query) and untreated lesioned (reference) cartilage explants. **(B)** Boxplots for top FDR-significant genes from RNA-seq as indicated (individual donor values are shown, color-coded by sample-ID; P: preserved; L: lesioned; T: lesioned treated (hiMSC+gel); FDR: false discovery rate; VST: normalized, variance-stabilized counts for mRNA reads). **(C)** Boxplots for genes as indicated, relevant for cartilage homeostasis. **(D)** Visualization of significant pathway enrichment among nominal significant genes with inverse direction of effect in comparison to FDR-significant genes identified with RNA-seq for preserved and lesioned cartilage (*22*)

### OA pathophysiological changes in chondrocyte signaling in human ex vivo cartilage explant models in response to hiMSC-treatment and OA responsive genes

Next to transcriptome wide analyses, we sought to confirm changes in chondrocyte signaling in response to hiMSC+gel treatment that mark cartilage integrity (*TNFRSF11B*) as well as chondrocyte phenotypic states such as *COL2A1, FRZB, SOX9* (anabolic healthy state) and *MMP13*, and *ADAMTS5* (catabolic osteoarthritic state) and plotted VST-normalized gene expression level in preserved, lesioned, and lesioned treated (hiMSC+gel) human cartilage explants. As shown in **Figure 3C** we observed that hiMSC+gel treatment generally restored gene expression towards the preserved cartilage level, albeit significant only for *COL2A1* (FC=2.1, *P*=1.6×10^−2^). It is noteworthy to mention that bulk RNA-seq data is per definition reporting on tissue wide changes.

Furthermore, we investigated to what extend the hiMSC+gel treatment reverses transcriptome wide markers of OA pathophysiology. In doing so, we overlayed hiMSC DEGs (DEG_hiMSC, *P*≤0.05) with those previously identified with OA pathophysiology (RAAK study DEG_OA, FDR<0.05)(*22*) and prioritized genes that showed an inverse effect, i.e. genes marking a restoring effect upon hiMSCs treatment towards preserved macroscopically normal cartilage (n=92, **Table 1**). To visually explore these genes, we generated a protein–protein interaction network among these 92 genes showing a highly significant gene enrichment for genes acting in the cellular Zinc ion homeostasis pathway (*P*=4.2×10^−4^), represented by genes such as *SLC30A1* and *SLC39A14* encoding respectively an Zn2+ exporter, and importer but also *MT1F, MT1G*, and *MT1H* encoding metal-ion binding proteins including Zn2+ ions (**Fig. 3D** and **Fig. S2**). Moreover, a significant local network cluster was marked by genes involved in collagen formation and cell adhesion (*P=*2.5×10^-2^), such as *ITGA5*, and *COL9A2*. These results suggested that the shift in sterol and cholesterol syntheses upon hiMSCs+gel treatment, is concomitant with an alleviation of the OA pathophysiological cellular Zinc-ion imbalances. Notably the catabolic cascade of Zinc-ion imbalances in OA pathophysiology is marked by *MMP3, MMP13*, and *ADAMTS5*.(*32*)

**Table 1.**
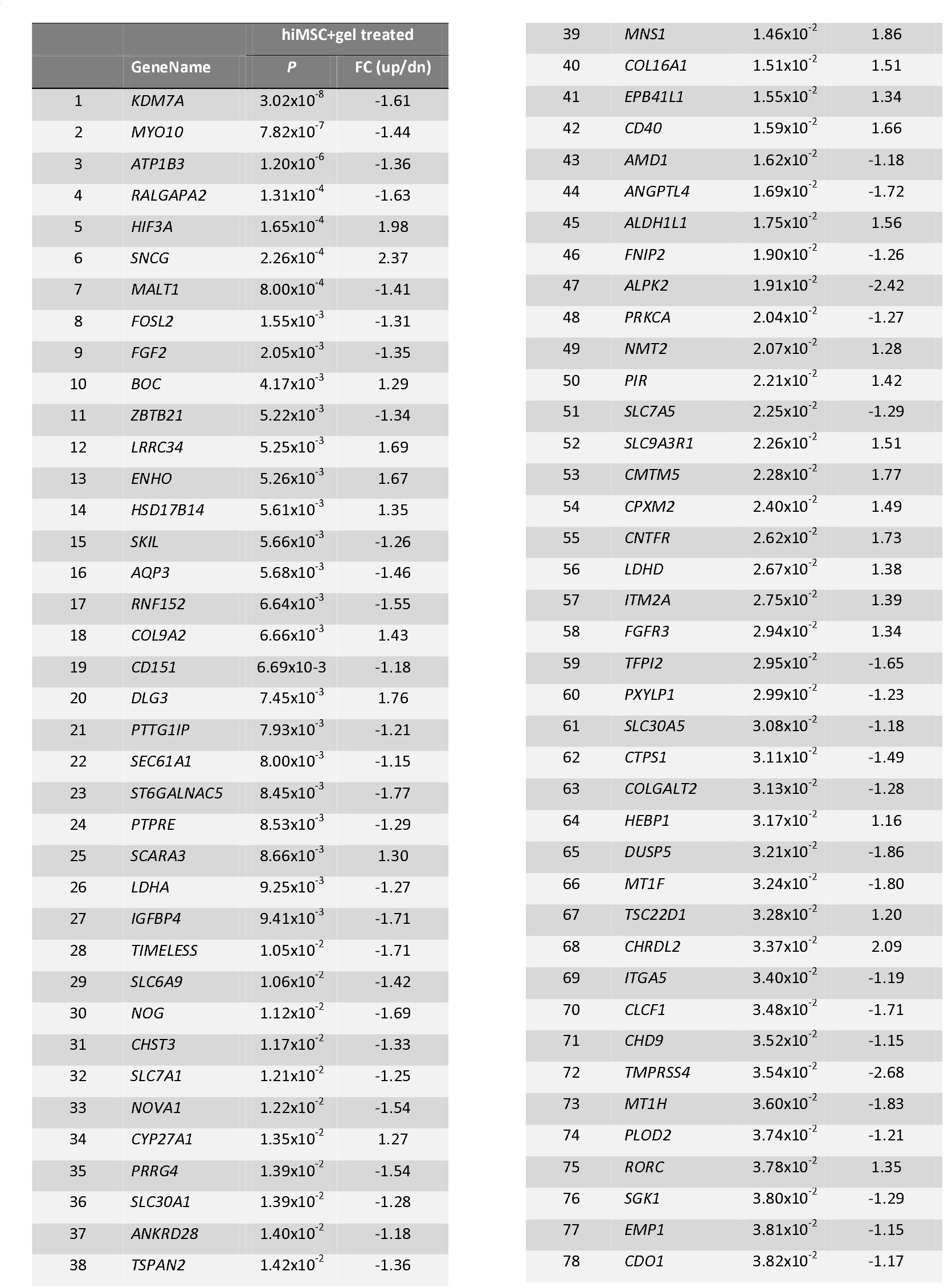
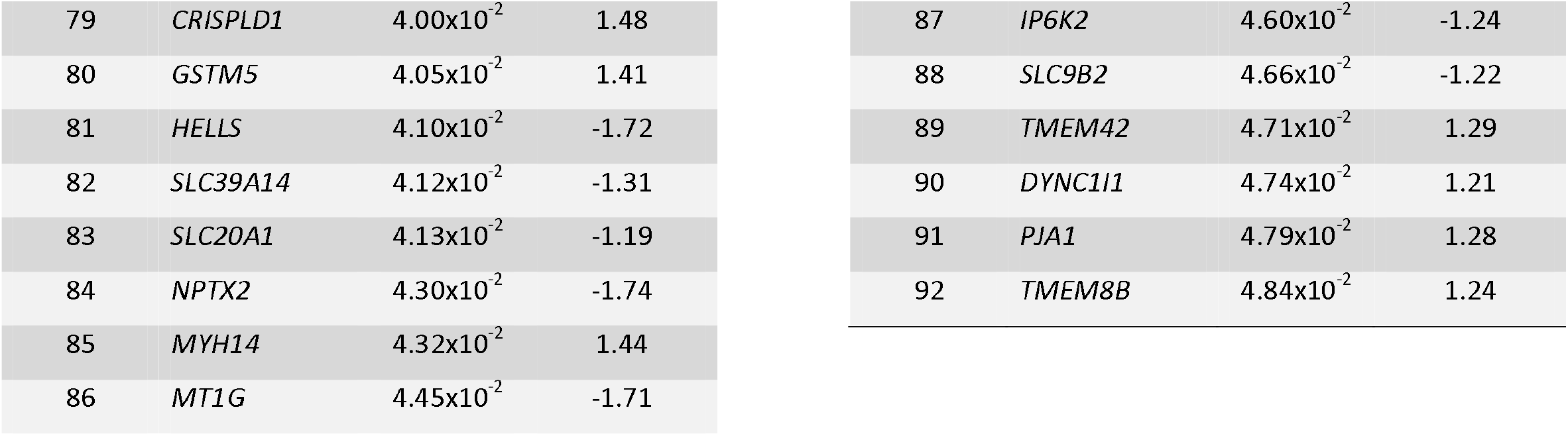
n=92 DEGs with hiMSCs treatment (*P*≤0.05) that have an inverse fold change (FC) with that of the previously found DEGs with OA pathophysiology (FDR≤0.05).(*22*)

### Direct cellular mode-of-action of hiMSC treatment on human chondrocyte cell populations by spatial transcriptomics analysis

Finally, we aimed to identify the mode-of-action of hiMSC+gel treatment on individual chondrocyte cell populations within native human cartilage tissue while preserving their *in situ* context i.e. providing information across articular cartilage layers. To this end, we performed spatial transcriptomics (Xenium, 10X-Genomics) with a custom probe panel of 295 genes (**Data file S3**) that included common cartilage and bone markers, as well as genes marking cholesterol and sterol metabolism and cellular Zinc ion homeostasis pathways, identified by the bulk RNA-seq. Spatial transcriptomics was performed on (n=7) untreated (control) and treated lesioned articular cartilage explants from two donors of the RAAK study (**Fig. S3**, donor-2: one untreated and two treated cartilage explants; donor-6: one untreated and three treated cartilage explants).

In total 22,669 cells were detected. After quality control and removal of cells with fewer than 10 transcripts, 3,853 (donor-2) and 3,465 (donor-6) high quality cells remained for further analysis. UMAP dimensionality reduction, followed by Leiden clustering algorithm were applied to each donor separately. We identified 10 cell clusters in donor-2 (**Fig. 4A** clusters 0–9) and 9 cell clusters in donor-6 (**Fig. S4A** clusters 0–8). This means that among the lesioned and hiMSC-treated native human articular cartilage tissues, similar but distinct chondrocyte cell populations exist. To capture cellular characteristics of these chondrocyte populations, we next plotted top 5 differentially expressed genes per cell cluster, by comparing each cluster against all others. As shown in **Figure 4B** and **Figure S4B**, we observed a similar and exclusive gene expression pattern for cell cluster-9 in donor-2 and cluster-4 in donor-6 characterized by expression of *DMP1*, encoding dentin matrix acidic phosphoprotein 1, a well-known gene expressed in osteocytes. On the other hand, we showed that almost all cell clusters had high marker gene expression of the cartilage proteoglycan *ACAN* (**Fig. 4C, Fig. S4C**).

**Fig. 4.**
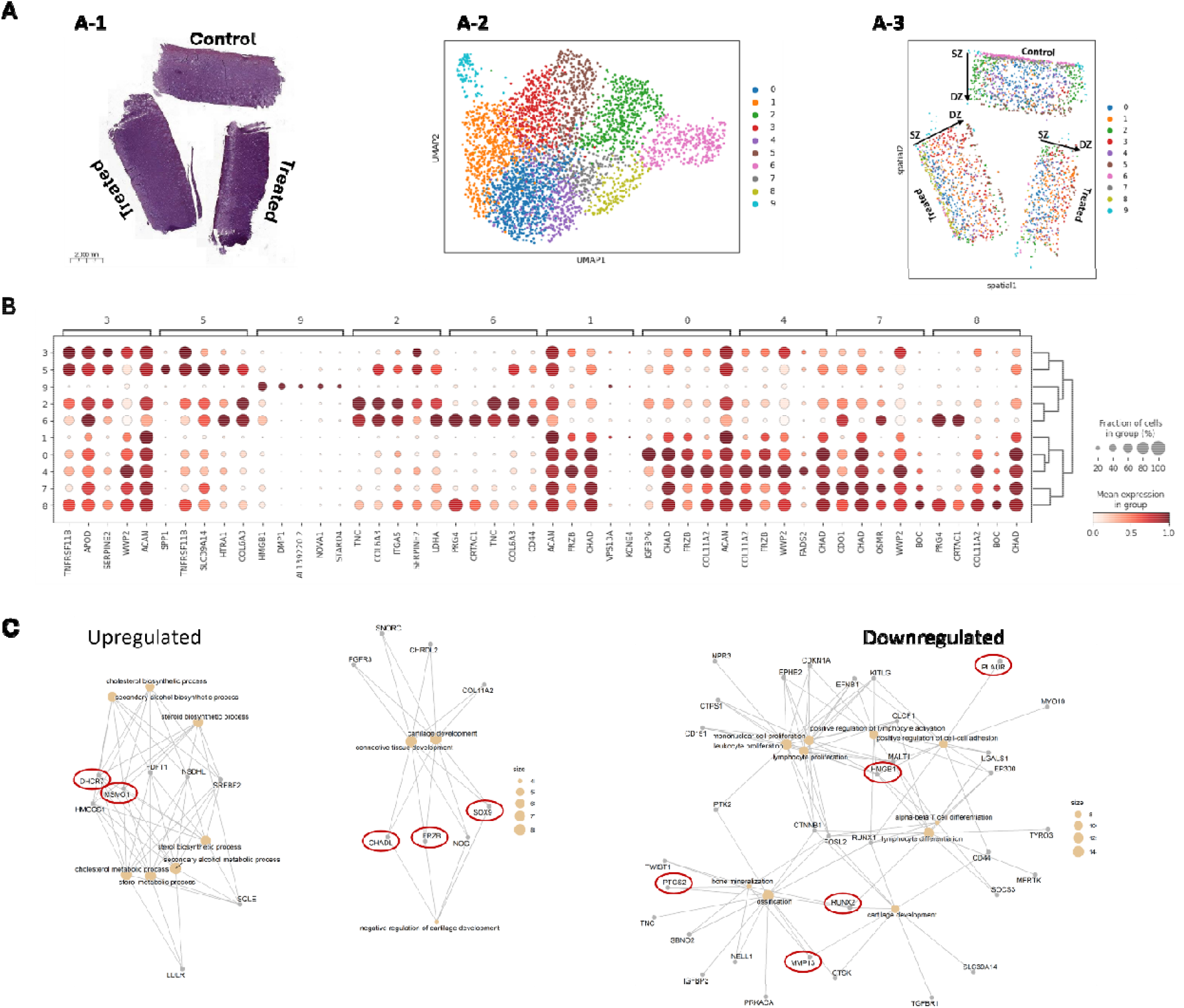
Spatial transcriptomic profiling of OA cartilage explants treated with hiMSC-gel. Spatial transcriptomics of paired OA-lesioned cartilage explants of donor-2. **(A)** Representative explants from donor-2: (A-1) untreated (control) and treated (hiMSC+gel) explants. (A-2) UMAP analysis identified 10 transcriptionally distinct clusters. (A-3) Clusters remapped to their spatial coordinates within the explant. **(B)** Dot plot showing differentially expressed genes (DEGs) across clusters. **(C)** Visualization of significant pathways enriched among differential upregulated (left panel) and downregulated (right panel) genes between treated cluster-8 cells as query and control cluster-6 as reference while showing links between genes and biological processes.

Upon mapping the cell clusters back to their *in situ* location (**Fig. 4A-3, Fig. S4A-3**), we revealed that the chondrocyte cell population with the exclusive gene expression pattern (DMP1, cell cluster-9 in donor-2 and cluster-4 in donor-6) mapped to the edges of control *ex vivo* explants, likely representing a cut artifact. More compelling, we identified in donor-2, exclusive cell populations in the superficial zone (SZ) of untreated (control, cluster-6, pink cells) and hiMSC-treated (treated, cluster-8, green cells) lesioned OA cartilage explants; i.e. SZ cell populations stratified by treatment and characterized by a unique gene expression profile (**Fig. 4A-3, Fig. 4B**). Similarly for donor-6, distinct cell clusters in the SZ layer were identified in control (cluster-3) and treated samples (cluster-1), although the SZ was less well defined due to the more severe damage (**Fig. S4A-3, Fig. S4B**). We reasoned that the gene expression profile of these the unique superficial cell clusters in control and treated samples could be explained the action of hiMSC+gel treatment.

To identify the mode-of-action of hiMSC+gel treatment specifically for the SZ cell populations in lesioned OA cartilage in donor-2, we performed differential gene expression analyses (control cluster-6 versus treated cluster-8) followed by gene enrichment analyses for biological process. In **Figure 4C**, we plotted enriched biological processes among significant differential upregulated (left panel) and downregulated (right panel) genes. Among the upregulated genes in the SZ of hiMSC-treated chondrocytes (**Fig. 4C** left panel), we confirmed sterol (FDR=9.9×10^-26^) and cholesterol (FDR=1.8×10^-23^) pathway genes *MSMO1, FADS1*, and *DHCR7*. Even more compelling, was the upregulation of well-known cartilage genes such as *SOX9, FRZB, ACAN*, and *CHAD*, all involved in connective tissue and cartilage development. Regarding the downregulated genes (**Fig. 4C**, right panel) we recognized among others, catabolic genes *MMP13, ADAMTS5, PTGS2* but also ossification pathway genes *RUNX2, DIO2, TNC*, and genes that act in senescence pathways such as *HMGB, PLAUR*, and *SERPINE1* (**Data file S7**). Notably in this respect was that changes in *TNFRSF11B (*Mankin-score) and cellular Zinc ion homeostasis pathways (OA responsive markers to hiMSC treatment) were not identified in the DEG genes between SZ cartilage layers of treated and untreated explants. Upon a closer examination, we identified that the cluster-1 cell population (orange cells) that become prominent in the middle and deep cartilage zone of hiMSC treatment, are characterized relative to all other cell populations by low expression of *TNFRSF11B* (logFC=-2.25, FDR=1.0×10^-73^) associated to Mankin-scores, low expression of *SLC30A1* (logFC=-1.63, FDR=5.7×10^-43^), and *SLC39A14* (logFC=-1.69, FDR=2.05×10^-56^) associated to OA related Zinc-transporters, but also *MT1H* (logFC=-0.68, FDR=3.1×10^-4^), *MT1G* (logFC=-0.43, FDR=2.0×10^-7^) and *MT1F* (logFC=-0.60, FDR=2.0×10^-2^) associated to OA related metal-ion binding proteins. Even though cartilage layers in donor-6 were less well defined, similar genes and pathway changes were identified upon hiMSC+gel treatment (**Fig. S4C**).

To identify the most consistent treatment-evoked transcriptional changes across donors, we also performed a fixed-effects meta-analysis, focusing on the SZ-clusters again. As shown in **Data file S8**, meta-analysis identified 126 significantly differentially expressed genes (FDR≤0.05), including 38 up- and 88 downregulated genes. Visualization of the gene-interaction network (**Fig. S5**) of upregulated genes in the SZ-cluster confirmed enrichment for biological processes related to cartilage development featuring *CHADL, FRZB, SOX9*, and sterol and cholesterol pathways featuring *MSMO1, DHCR7*, and *EBP*. Notably in the meta-analyses was the appearance of *NGF* to be significantly downregulation in the SZ-clusters. Since *NGF* encodes the well-known OA pain associated nerve growth factor, this result suggested an attenuation of neurogenic and pain-related pathways within the hiMSC+gel treatment in cartilage.

Together the ability to sensitively map spatially defined molecular profiles of individual chondrocyte cell populations revealed that hiMSC+gel treatment, restored the molecular identity of osteoarthritic chondrocytes toward a healthy articular cartilage cellular phenotype, particularly in the superficial cartilage layer.

## DISCUSSION

Targeting articular cartilage degeneration remains a central challenge in OA, with limited disease-modifying therapies available. In the current study, we explored therapeutic efficacy of human induced pluripotent-derived therapeutic stem cells (hiMSCs) across *in vivo* mouse and *ex vivo* human osteoarthritis models. We showed that hiMSC treatment in a DMM-mouse model of OA significantly reduced OARSI damage scores, which was affirmed by a decrease in the catabolic marker Mmp13 and an increase in the anabolic marker Col2. These treatment effects appeared, irrespective of potential modifying factors such as xeno-free media or thermosensitive hydrogel carrier. Even more, the thermosensitive hydrogel improved injectability and showed an independent beneficial effect. Subsequently treatment of hiMSC+gel in the human osteoarthritic cartilage explant model, showed a transcriptome-wide significant activation of the cholesterol and sterol biosynthesis pathways marked by differential expression of genes such as *MVD, DHCR7, MSMO1, FABP3*. Additionally, we showed that these changes alleviated OA associated imbalances in the cellular Zinc-ion homeostasis pathways, represented by genes such as *SLC30A1, SLC39A14* and *MT1F, MT1G, MT1H*. Spatial transcriptomics then sensitively captured that hiMSC+gel treatment evoked, specifically at the superficial cartilage layer, a consistent upregulation of healthy chondrocyte markers such as *CHAD, ACAN, FRZB*, and *SOX9*, alongside a suppression of catabolic and inflammatory mediators such as *SERPINE1, SPP1, MMP13*, and *ADAMTS5*. Together our study highlighted that hiPSC-derived stem cell therapy using hiMSCs restored the molecular identity of osteoarthritic chondrocytes. Being hiPSC-derived, our therapeutic cells support a scalable ‘off-the-shelf’ solution to treat osteoarthritis, in the near future.

Our study showed that hiMSCs+gel activated sterol/cholesterol biosynthesis, concomitant with an alleviation of the OA pathophysiological cellular Zinc-ion imbalances (*MT1F, MT1G, MT1H* and *SLC30A1, SLC39A14*). Subsequent spatial transcriptomics, indeed revealed downregulation of the associated catabolic cascade of Zinc-ion imbalances (*MMP13* and *ADAMTS5*).(*32*) Regarding the sterol biosynthesis pathway it is worthy to not that *DHCR7* is essential for converting 7-dehydrocholesterol to cholesterol, a step critical for proper primary cilium formation and Hedgehog signaling. Since primary cilia act as important sensory organelles in chondrocytes, regulating responses to mechanical and biochemical signals that maintain cartilage homeostasis.(*33, 34*) Similarly, *MSMO1* encodes methyl sterol monooxygenase 1, an enzyme involved in the demethylation of sterol intermediates during cholesterol biosynthesis, essential for proper chondrocyte differentiation, growth plate formation and endochondral ossification.(*35*) These findings raise the possibility that reactivation of endogenous sterol / cholesterol biosynthesis represents a previously underappreciated mechanism of cartilage repair (*30*), rather than reflecting pathological lipid accumulation. (*36*) We advocate, therefore, that induction of the canonical sterol biosynthetic machinery by hiMSC treatment, may represent a regenerative adaptation that restores the metabolic capacity of chondrocytes required for matrix-producing anabolic activity.(*37*) The concomitant normalization of OA-associated zinc homeostasis, particularly across the deeper zones suggests that this metabolic reprogramming occurs alongside attenuation of the catabolic cascade characteristic of osteoarthritic chondrocytes.(*32*)

In our study, we employed both an *in vivo* DMM mouse and a human *ex vivo* OA cartilage explant model and assessed the therapeutic efficacy of hiMSCs. The preclinical *in vivo* DMM model, a widely used *in vivo* model to study OA(*20*), allowed us to study the hiMSCs treatment effect for the entire synovial compartment of the joint, hence allowing inflammatory processes to be taken along in disease progression.(*20*) It also enabled us to test the effect of recently developed thermosensitive hydrogel for i.a. injections in a controlled joint environment.(*14*) In principle, however, the DMM model reflects a post-traumatic OA phenotype that is per definition not equivalent to human OA, which is multifactorial and involves age-related factors. Also in our DMM model we used only males whereas, animal studies inherently conflict with the principles of the 3Rs (replacement, reduction, refinement(*38*)). Therefore, we here introduced human cartilage explant model providing a faithful representation of human lesioned OA cartilage, directly obtained from male and female patients that underwent a joint replacement surgery due to OA.(*30*) Notably, despite donor heterogeneity, our hiMSC+gel treatment revealed upregulation of genes involved in cholesterol biosynthesis, whereas spatial transcriptomics revealed consistent changes in chondrocyte populations across donors. Together, we leveraged the complementary strengths of physiological disease complexity and direct human tissue relevance, providing compelling evidence that the identified mechanism of action is both biologically robust and clinically translatable.

Our findings link therapeutic outcomes of hiMSC treatment to precise spatially resolved molecular changes in human tissue, that would otherwise be obscured by heterogeneous cell populations. As such, spatial transcriptomics sensitively unravelled that the hiMSC+gel treatment, evoked restoration of a healthy chondrocyte identity (*SOX9, FRZB, CHAD, COL2A1*) specifically in superficial cartilage layers. An OA affected cartilage layer that is notoriously depleted of anabolic activity.(*39*). *SOX9*, for that matter, is a regulator of chondrogenesis and essential for the activation of *COL2A1* and the maintenance of healthy chondrocyte phenotypic state. Notably in this respect is the observation that targeted genes had a significant enrichment for CCAAT-box sequences e.g. found in the proximal core promoter of *SOX9*(*40*) that is recognized by the trimeric NF-Y transcription complex.(*41, 42*) Moreover, *FRZB*, an antagonist of Wnt signaling, is known to protect cartilage homeostasis by inhibiting hypertrophic differentiation and osteophyte formation.(*43*) *CHADL* is involved in ECM organization and collagen binding, supporting proper cartilage architecture and is a negative modulator of chondrocyte differentiation.(*44*)

On a different note, hiMSC+gel treated superficial chondrocyte cell populations revealed significant reduced expression of *NGF*, a gene that particularly acts in the biological processes of axon development. Since this pathway is strongly associated with both OA associated pain sensitization and neurogenic inflammation (*45*), we advocate that hiMSC treatment could also contribute to OA pain management. Nonetheless, a perceived weakness is that we did not include pain-related behavior in the *in vivo* experiment. Another limitation of the performed animal model was that due to limited number of mice, joint tissue damage resulting from DMM was not determined prior to treatment. As such, we cannot confirm whether the cells only slow-down the development of OA or whether it can really restore the damage. Results of the human *ex vivo* OA cartilage explants across OA donors, however, did convincingly demonstrate a restoration of the OA chondrocytes toward a healthy chondrocyte state, likely able to initiate cartilage repair. This was further supported by the observed changes in TNFRSF11B/OPG expression that strongly correlates to cartilage damage measured by the Mankin-score.

In conclusion, by combining an *in vivo* animal and introducing a human OA cartilage explant model, we position treatment with hiMSCs as a potent ‘off-the-shelf’ disease modifying therapy for OA. Our multitiered approach allowed linking of therapeutic outcomes of hiMSC treatment to precise spatially resolved molecular changes in human tissue, that would otherwise be obscured by heterogeneous cell populations. In doing so we demonstrated that treatment with hiMSC+gel restored the molecular identity of osteoarthritic chondrocytes toward a healthy state. The insights provided by our work contribute to clinical development and optimization of next-generation regenerative stem cell therapies for cartilage disorders. In this respect, we envision that our here introduced human lesioned OA cartilage explant model, could facilitate selection of potent therapeutic cells prior to cryopreservation and cell banking at large scale.

## Supporting information

SupplementaryMaterial_hiMSCSpecial

## List of Supplementary materials

- Supplementary Materials and Methods
- Fig. S1 to Fig. S4
- Data file S1 to S8 (Excel files)
- References only cited in Supplementary materials: none

## Acknowledgements

We thank all study participants of the RAAK and the GARP study, both supported by the Leiden University Medical Centre.

## Contributions

Yolande FM Ramos: Conception and design, collection and/or assembly of data (ex vivo experiments), data analysis and interpretation, final approval of manuscript.

Sana S. Sayedipour: Collection and/or assembly of data (in vivo experiments), data analysis and interpretation, manuscript writing, final approval of manuscript.

Georgina Shaw: Collection and/or assembly of data, final approval of manuscript.

Margo Tuerlings: Collection and/or assembly of data, final approval of manuscript.

Timo Schomann: Collection and/or assembly of data, final approval of manuscript.

Eka Suchiman: Collection and/or assembly of data, final approval of manuscript.

Rachid Mahdad: Collection and/or assembly of data, final approval of manuscript.

Hailiang Mei: Collection and/or assembly of data, final approval of manuscript.

Davy Cats: Collection and/or assembly of data, final approval of manuscript.

Luis J. Cruz: Collection and/or assembly of data, final approval of manuscript.

Mary Murphy: Collection and/or assembly of data, final approval of manuscript.

Ingrid Meulenbelt: Collection and/or assembly of data (in vivo experiments), data analysis and interpretation, manuscript writing, final approval of manuscript.

## Data availability statement

All data are available upon reasonable request. RNA-seq dataset for human articular cartilage explants from the RAAK study have been deposited at the European Genome-phenome Archive (<u>EGA</u>) under number EGAD50000000570.

## Ethics approval

The project “Dose finding & potential adverse effects observations in the DMM model” was conducted at the Leiden University Medical Center and was approved by the Animal Welfare Committee (IvD) under number AVD1160020171405-PE.18.101.006 on Apr 21, 2023.

The Medical Ethics Committee of the LUMC gave approval for generation of hiPSCs from skin fibroblasts of healthy donors under number P13.080 on July 2 2014 (Parapluprotocol: hiPSC). Informed consent was obtained from all donors.

Collection of hBMSCs from OA patients undergoing total joint replacement surgery is approved by the Medical Ethical Committee of the LUMC within the ongoing RAAK study and available under numbers P08.239 and P19.013.

The authors declare that they have not used AI-generated work in this manuscript.

