## SupplementaryMaterial_hiMSCSpecial for "Spatial transcriptomics defines the mechanisms of hiPSC-derived stem cell-mediated repair in human articular cartilage": 260716_Spatial_SupplementaryMaterial.pdf

### Supplementary Materials and Methods:

#### *In vivo osteoarthritis model*

Twelve week old male C57BL/6J mice were purchased from Charles River Laboratories (Charles River, Chatillon-sur-Chalaronne, France). Animal procedures were all conducted at the Leiden University Medical Center and were approved by the Animal Welfare Committee (IvD) under number AVD1160020171405-PE.18.101.005. All mice were housed in groups in polypropylene cages on a 12-hour light/dark cycle with unrestricted access to standard mouse food and water. Knee OA was induced in the mice by surgical destabilization of the medial meniscus (DMM) on the right knee joint (N=42) as described previously.(*25*) Briefly, 30 minutes before surgery, pre-operative analgesia (Buprenorphine) was administered by subcutaneous injection and the animal was then placed under isoflurane anaesthesia. Once the animal no longer displayed reflexes while retaining constant breathing, a 1-cm longitudinal medial para-patellar incision was made on the right knee to gently expose the joint. The tibial plateau was then cut upon lateral dislocation of the patella and patellar ligament. After transection, the knee joint capsule was closed with a 6-0 absorbable suture and the skin closed with biological glue. Mice were immediately transferred to a heated post-operative recovery room. Post-surgery, all animals received buprenorphine HCl (Vetergesic; Alstoe Animal Health, York, UK) sub-cutaneously every 8hr for 48hr and they were monitored daily to ensure good health.

To investigate the therapeutic efficacy of hiMSCs and hBMSCs cultured in medium containing fetal calf serum (FCS) or in a chemically defined and xeno-free medium (PurStem (PS)), Three weeks after the surgery we performed intra-articular (i.a.) injections (30G needle). DMM mice were treated with a one-time i.a. injection of 6µL PBS, either or not with hydrogel and $1x10^6$ hiMSCs cultured in PS medium and/or hydrogel, $1x10^6$ hiMSCs cultured in serum containing medium and/or hydrogel, $1x10^6$ hBMSCs cultured in PS medium with hydrogel, and $1x10^6$ hBMSCs cultured in serum containing medium with hydrogel. Animals were randomly allocated to the experimental groups (n=6-12) using a dice. Moreover, to ensure blinded treatments, i.a. injections were performed by a colleague not involved in randomization while the injection samples were prepared and coded by another colleague not involved in the animal experiment. Mice were monitored daily to confirm general health indicators according to their body weight and knee diameter, and were sacrificed 35 days after the injection for analysis.

#### *Histological assessment mouse knee joints*

For histological assessment of *in vivo* samples, we followed the OARSI guidelines.(*25*) The knee joints were fixed in 4% paraformaldehyde for 24hr, followed by 3 weeks decalcification in 10% EDTA (pH 7.4) at room temperature. They were then embedded in paraffin and knee joints were cut into 5 µm sections. The sections were stained with Safranin-O/Fast green and Haematoxylin & Eosin Y (H&E), and examined by light microscopy to evaluate the cartilage damage of the femur and tibia in knee joint. The scoring was performed by three independent researchers who were blinded to the conditions and to the scores of the other investigator. The results were averaged and the OA score of the sections was taken as the representative score of the knee joint, as described elsewhere.(*39, 40*) Immunohistochemistry staining was performed on knee joint sections for collagen type 2 (Col2) and Mmp13. For both staining procedures, slides were blocked for endogenous peroxidase using 0.3% $H_2O_2$ for 10min at room temperature. Antigen retrieval was performed with 25µg/mL Proteinase K (Prot K) prepared in 0.1M Tris/HCL, pH 5.0 for 10min at 37°C, followed by 30min treatment with hyaluronidase (5mg/mL in Tris/HCL pH 5.0). For both types of staining, all slides were blocked in 5% PBS-BSA for 30min at room temperature. Primary Col2 antibody incubation was done overnight at 4°C with 0.2µg/mL Col2 mouse monoclonal antibody (sc7271, Santa Cruz Biotechnology, Santa Cruz, Dallas, TX, USA) or with 0.2µg/mL normal mouse IgG1 as isotype control (sc3877, Santa Cruz, Dallas, TX, USA). Primary Mmp13 antibody incubation was done overnight at 4°C with 2µg/mL Mmp13 monoclonal antibody (sc515284, Santa Cruz Biotechnology, Santa Cruz, Dallas, TX, USA) or with 2µg/mL normal mouse IgG1 as isotype control (sc3877, Santa Cruz, Dallas, TX, USA). All antibodies were diluted in 5% PBS-BSA. The next day, slides were incubated with anti-mouse HRP (Envision, Dako, CA, USA) for 30min at room temperature and subsequently incubated with liquid DAB + 2-component system (Agilent, Santa Clara, CA, USA) for 5min. Sections were counterstained with hematoxylin, dehydrated, cleared in xylene and cover slipped with Eukitt® mounting medium (Sigma-Aldrich, Saint Louis, USA) and microscopy images were quantified using ImageJ. Hereto, color channels were split and selected the red channel for further processing. Next, using a rolling ball algorithm the background noise was removed and the sample was segmented. Within this segmentation of the sample, we calculated the average staining intensity.(*41*)

***RNA-sequencing***

RNA-seq analyses was performed on controls (n=7 preserved, n=12 lesioned OA cartilage samples from N=9 donors) and treated samples (n=17 lesioned OA cartilage samples treated with hiMSCs in the thermosensitive hydrogel from N=9 donors). Hereto, RNA was extracted with chloroform and purified using the RNeasy Mini Kit (QIAGEN). RNA libraries were generated (polyA enriched), and three-prime RNA sequencing was performed using Illumina NovaSeq 6000 according to the standard operating procedures based on the Illumina protocol for Paired-End Sequencing (PE150bp) with v1.5chemistry. Subsequently, the in-house available pipeline BioWDL (https://biowdl.github.io/RNA-seq/v5.0.0/index.html), developed by the Sequencing Analysis Support Core at Leiden University Medical Center, was used to process FASTQ files including adapter clipping with cutadapt.v2.10, and QC with FastQC.v0.11.9 and MultiQCv. 1.9. Furthermore, mapping was performed with STAR.v2.7.5a software and expression quantification and transcript assembly using HTSeq-Count.v.0.12.4. The reads were aligned to the human reference genome GRCh38 and Ensembl gene annotation version 109, and RNA-seq data was normalized using the DESeq2_v.1.30.0 R package. Before analysis, the data was transformed using the variance-stabilizing transforming (VST) method. Differential expression analysis was performed using the DESeq2 package v. 1.30.0 using R version 4.0.2. A general linear model (GLM), assuming a negative binomial distribution was applied, followed by a Wald-test to compare the control with treated explants. Benjamini-Hochberg multiple testing corrected *P*-values with a significance cut-off of 0.05 are reported as False Discovery Rate (FDR).

### Legend Supplementary Figures

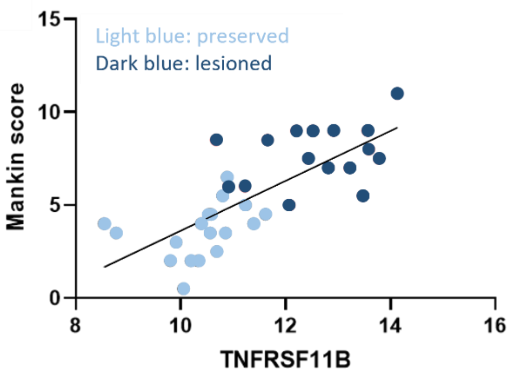


**Fig. S1. Correlation Mankin score and *TNFRSF11B* gene expression levels** (ρ=0.81 with *P*=8.6x10$^{-9}$).

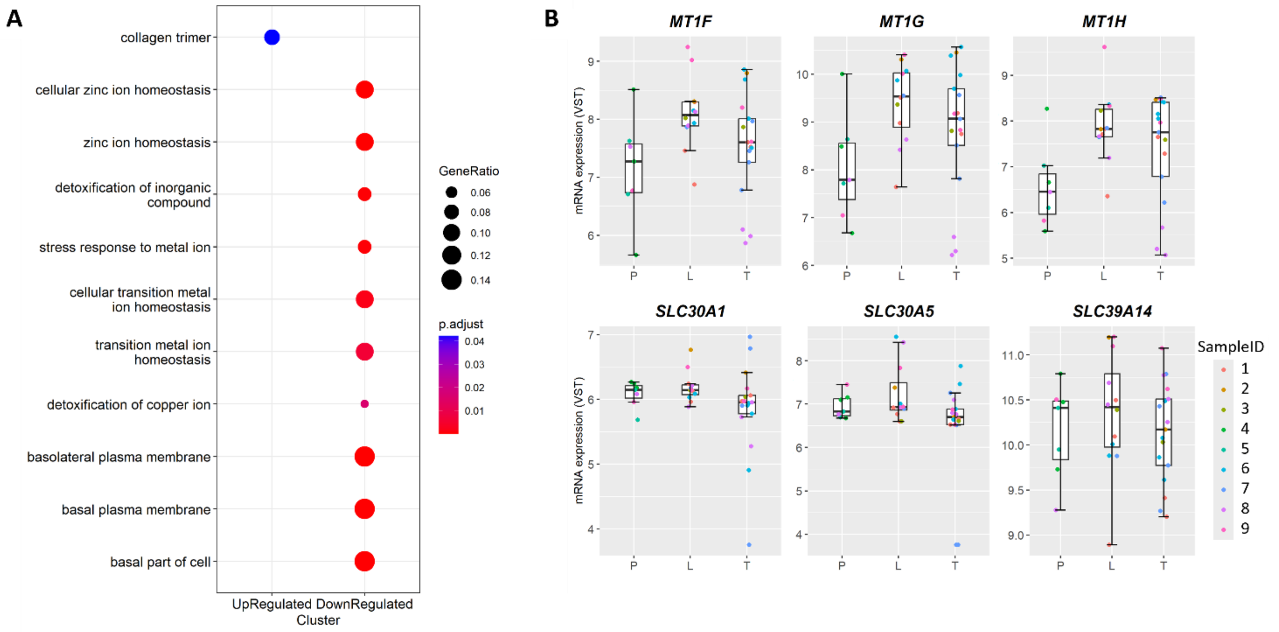


**Fig. S2. Pathways and boxplots. (A)** Pathways that were significantly enriched among the genes (n=92) that showed inverse direction of effects when compared to changes with OA pathophysiology.(17) **(B)** Boxplots for identified genes as indicated, involved in cellular Zinc-ion homeostasis pathways and with inverse effects in response to hiMSC+gel treatment of lesioned cartilage explants from OA patients undergoing joint replacement surgery.

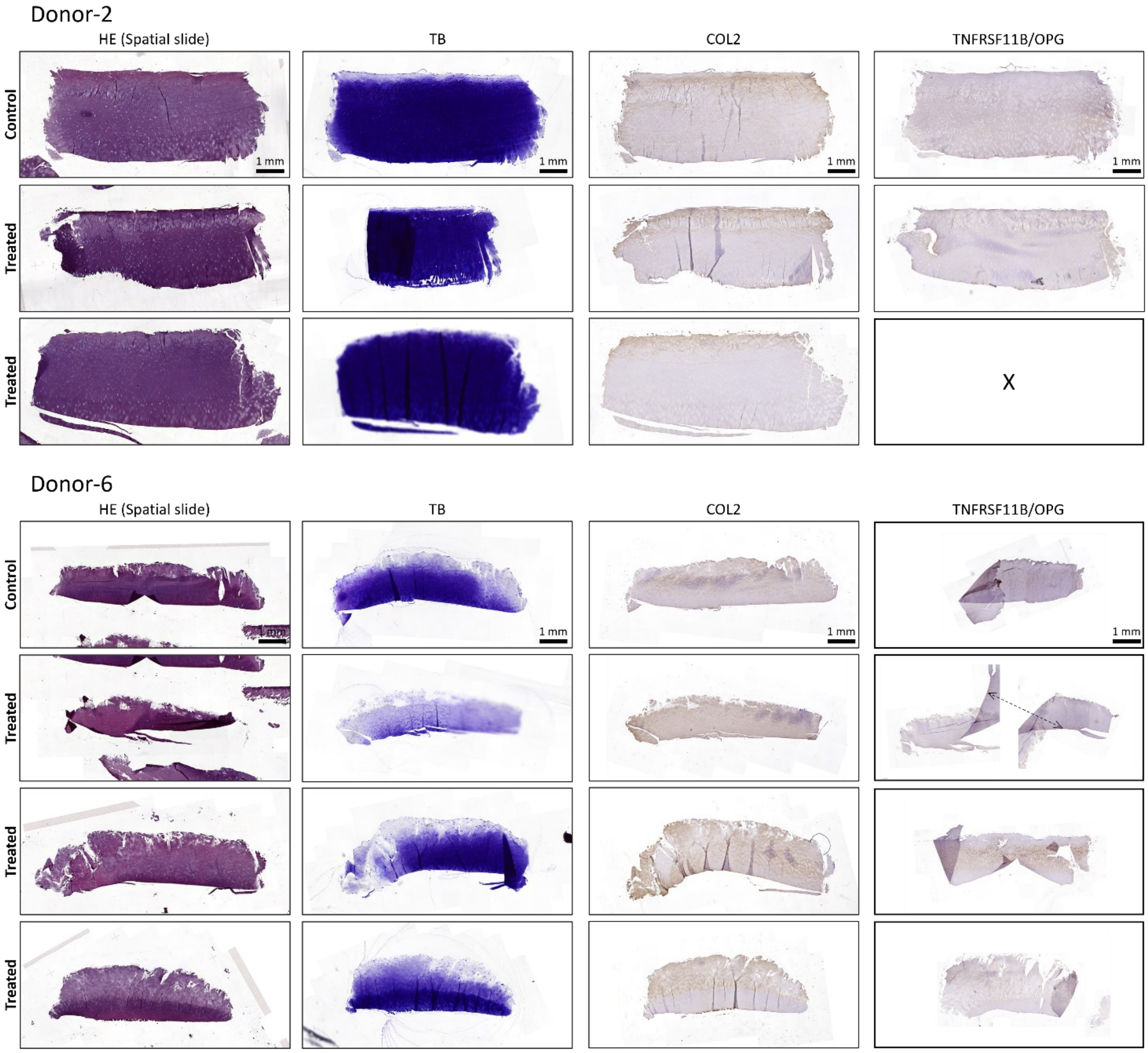


**Fig. S3. Representative images for explants included in spatial transcriptomic analysis.** Histology (HE and Toluidine Blue staining) and immunohistochemistry (COL2 and TNFRSF11B/OPG) for donor-2 and donor-6 (X section lost during procedure).

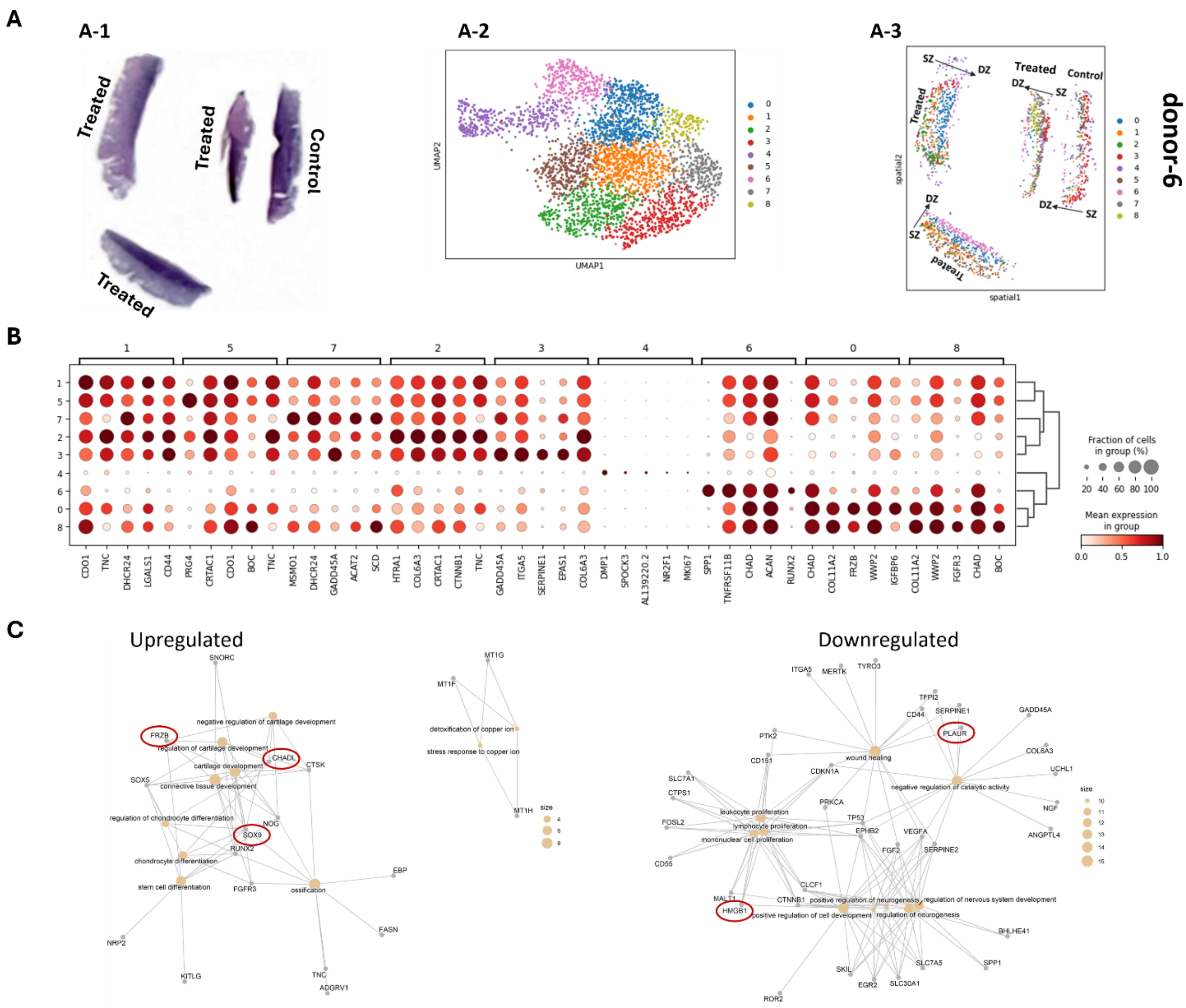


**Fig. S4. Spatial transcriptomic profiling of treated OA cartilage explants donor-6.** Spatial transcriptomics of paired OA-lesioned cartilage explants of donor-6. **(A)** Representative images for donor-6: **(A-1)** untreated control and hiMSC+gel-treated sections from spatial slides. **(A-2)** UMAP analysis identified 10 transcriptionally distinct clusters. **(A-3)** Clusters remapped to their spatial coordinates within the cartilage sections. **(B)** Dot plot showing differentially expressed genes (DEGs) across clusters. **(C)** Visualization of significant pathways enriched among differential upregulated (left panel) and downregulated (right panel) genes between treated cluster-3 cells as query and control cluster-1 as reference while showing links between genes and biological processes.

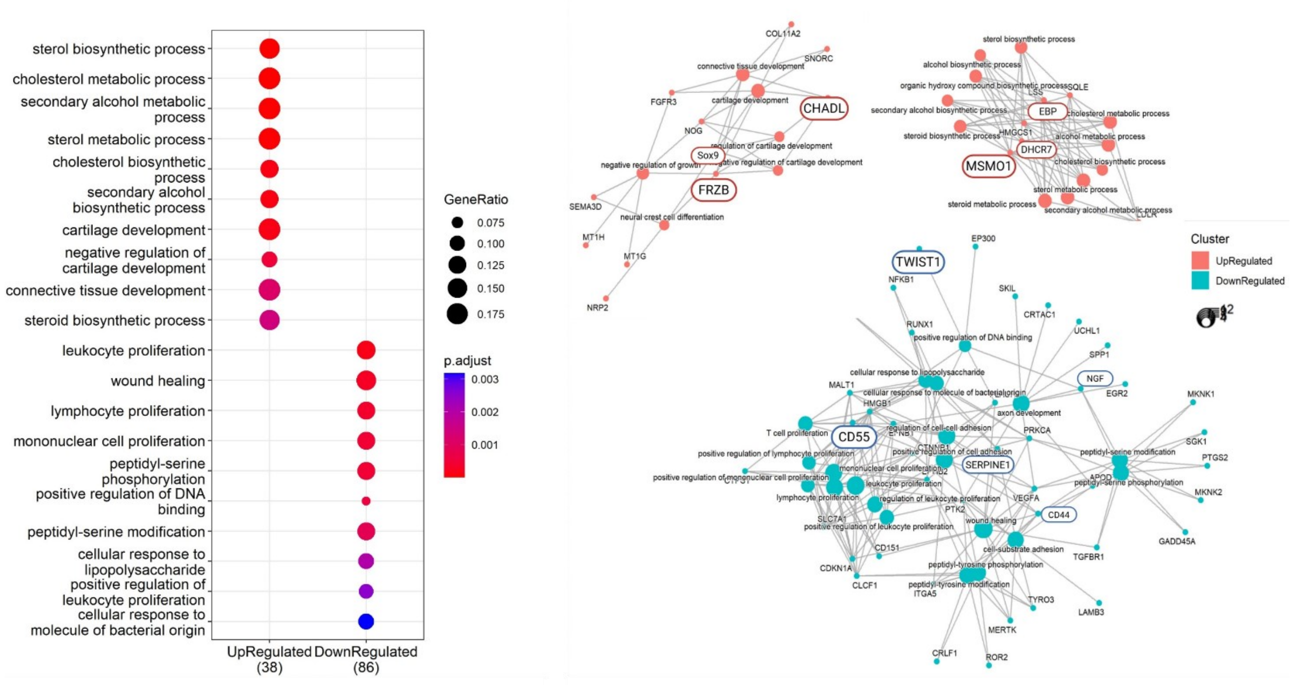


**Fig. S5. Pathways and protein-interaction networks that change in response to hiMSC+gel-treatment of lesioned articular cartilage from osteoarthritis patients as identified with spatial transcriptomics** (meta-analysis of genes expressed in superficial zone of donor-2 and donor-6).
